# Development and characterization of a biotechnological model suitable for studies of tumor cell extravasation and intravasation

**DOI:** 10.64898/2025.12.13.694106

**Authors:** Ekaterina V. Ivanovskaya, Georgii A. Bykov, Egor O. Osidak, Anastasia N. Sveshnikova

## Abstract

Metastatic dissemination remains the leading cause of mortality in malignant tumors, yet the processes of intravasation and extravasation of circulating tumor cells (CTC) are still not fully understood. Existing microfluidic experimental systems possess a number of limitations that prevent them from reproducing physiological conditions. Here we propose a perfused biotechnological system designed to model key stages of the metastatic cascade under controlled flow. The construction includes a parallel-plate flow chamber formed between polyethylene terephthalate plates and surrounded by a collagen gel containing life human dermal fibroblasts. Matrix parameters were optimized, and it was established that a collagen concentration of 20 mg/mL provides mechanical stability, sustained cell viability, and gel robustness under flow. The system also supports the formation of a two-component cellular microenvironment: fibroblasts embedded within the matrix and endothelial cells forming a layer on its surface. An integrated open reservoir effectively eliminated air bubbles and stabilized hydrodynamics, representing a major advantage over conventional microfluidic systems. 48 hour long perfusion of full medium with cells demonstrated long-term cell viability and preservation of channel geometry under continuous perfusion. The developed system combines the benefits of 3D hydrogels and dynamic models while overcoming critical limitations of classical microfluidic devices, and it may serve as a reproducible platform for studying mechanisms of metastasis.

## 1. Introduction

Metastasis remains the leading cause of cancer-related mortality, determining the prognosis for most patients even in the presence of advances in early diagnosis and local control of the primary tumor. The key yet least explored steps of the metastatic cascade are the intravasation and extravasation of tumor cells into and out of blood vessels. During intravasation, cells of the primary tumor penetrate through the endothelial barrier into the lumen of microvessels and enter the bloodstream. Extravasation enables circulating tumor cells (CTCs) to exit into a target tissue and initiate metastatic outgrowth. These processes largely define the aggressiveness of most solid tumors. However, the molecular and biophysical mechanisms underlying these stages remain insufficiently understood [1,2]. Their experimental reproduction is a complex biotechnological challenge due to the dynamic nature of the blood flow, the involvement of extracellular matrix and stromal cells, and the presence of gradients of mechanical and biochemical factors. This creates a strong demand for biotechnological models capable of recapitulating these biophysical conditions.

Classical two-dimensional models, 2D cultures, and Transwell systems provide basic information on cell migration and signaling, but they fail to reproduce the three-dimensional architecture of tumors, the mechanical properties of the matrix, and the influence of flow [3]. Therefore, in recent years, 3D models—such as spheroids, organoids, hydrogels, and bioprinted constructs—have been actively developed, offering a more accurate representation of tissue structure and the tumor microenvironment [4]. However, most of these 3D constructs remain static and do not reproduce key hydrodynamic parameters.

The next stage of development has been the emergence of “organ-on-a-chip” and “tumor-on-a-chip” technologies. These microfluidic models enable the integration of three-dimensional tissue structures with controlled perfusion. Recent reviews emphasize that such devices more accurately reproduce mechanical and chemical gradients than classical 2D/3D models and allow dynamic investigation of cellular behavior [5,6]. Within this field, specialized metastasis-on-a-chip platforms have been developed to analyze adhesion, transendothelial migration, and extravasation of tumor cells in real time [7].

Despite their advantages, microfluidic models of metastasis possess several fundamental limitations that become critical when studying intravasation and extravasation. Microchannels typically have a height of 50–150 μm, which significantly restricts the free migration of tumor cells, whose size often exceeds the diameter of capillaries. In microfluidic systems, passing through narrow channels inherently induces deformation-related stress, alters the mechanotype of cells, and may activate RhoA/ROCK signaling, an EMT-like phenotype, and a stress response uncharacteristic of native capillary venules [8]. Common materials used for microchips, primarily PDMS, are capable of absorbing cytokines, lipophilic ligands, and therapeutic molecules, complicating the modeling of native biochemical conditions and hindering quantitative interpretation of results [9]. The small thickness of the 3D matrix limits the ability to reproduce full mechanical and diffusion gradients typical of tumors [10]. Additionally, microfluidic systems are vulnerable to the formation of air bubbles, which complicates the maintenance of stable perfusion and introduces variability into experiments. Given these limitations, there is a growing need for alternative flow-based models that retain precise flow control while providing more physiological channel dimensions, resistance to degassing, compatibility with native fluids, and the ability to incorporate large, cell-laden 3D hydrogels.

The present study is dedicated to the development of a flow-based system and the optimization of its key parameters for investigating the intravasation and extravasation of tumor cells. The system consists of a channel formed between polyethylene terephthalate plates and surrounded by a collagen gel containing fibroblasts. A cell culture media or a suspension of tumor cells is continuously perfused through the channel. An additional open reservoir is used for bubble removal and stabilization of hydrodynamics. The aim of this work is to develop the system and optimize parameters such as gel concentration, incubation time, flow geometry, and a mathematical model of fluid movement within the channel of the flow chamber. Establishing such a system will expand experimental capabilities for studying key mechanisms of the metastatic cascade and help overcome limitations associated with microfluidic models.

## 2. Materials and Methods

### 2.1 Cell culture

Cell culture procedures were performed according to standard protocols described previously [11]. Briefly, endothelial cell line Ea.hy926 (ATCC, USA) and primary human dermal fibroblasts HDF (Koltsov Institute of Developmental Biology, Russian Academy of Sciences, passages 2–3) were cultured in 25 or 75 cm^2^ culture flasks (SPL Lifesciences, South Korea) in high-glucose DMEM medium (Servicebio, China) supplemented with 10% (v/v) fetal bovine serum (Himedia, India), 1% L-glutamine (PanEco, Russia), and 100 U/mL penicillin and 100 U/mL streptomycin (Thermo Fisher Scientific, USA). Upon reaching 80% confluence, cells were passaged using Versene solution (PanEco, Russia) and 0.05% trypsin (PanEco, Russia). The percentage of viable cells was assessed using a Goryaev chamber.

### 2.2 Preparation of collagen gel

Collagen gel was prepared under sterile conditions at concentrations of 10, 20, and 30 mg/mL by diluting the stock solution of type I collagen (Viscoll®, PA8, Imtek LLC, Russia) with complete DMEM medium. Mixing was carried out using two syringes connected by a connector, after which a suspension of human dermal fibroblasts (1×10^6^ cells/mL) was added to the gel (Fig. 1). The resulting gel was applied into the chamber or into the wells of a 6-well plate and allowed to polymerize in an incubator for 10–30 minutes depending on gel concentration. After polymerization, 1.5 mL of complete culture medium was added to the surface of the gel. Incubation was carried out for 3–5 days with regular monitoring using light microscopy. Final imaging was performed using light microscopy with staining with hematoxylin and eosin.

**Figure 1.**
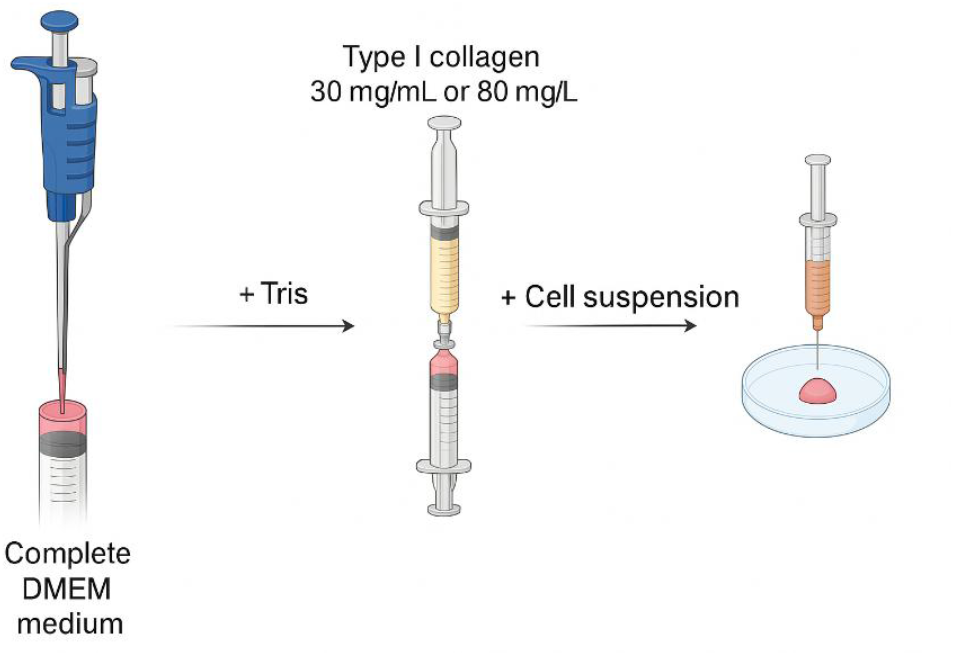
Schematic illustration of preparing collagen gel with embedded cells process.

### 2.3 Co-culture of multiple cell lines

To model tumor–stromal interactions, a co-culture of human dermal fibroblasts and the endothelial cell line Ea.hy926 was established. Fibroblasts were encapsulated in collagen gel (20 mg/mL) and incubated for 24 hours, after which Ea.hy926 cells were applied to the surface of the gel. Staining with hematoxylin and eosin was performed on days 5 and 8 of incubation.

### 2.4 Assembly of the chamber

The flow chamber consisted of a parallel-plate structure composed of three polyethylene terephthalate (PET) plates. The upper plate contained inlet and outlet openings positioned 30 mm apart; tubing was securely attached to these openings using hot-melt adhesive. The middle plate contained a central cavity for the application of collagen gel. The bottom plate was solid. All components were sterilized and the plates were joined using double-sided adhesive tape. The thickness of the internal channel between the upper and lower plates was 2 mm, ensuring flow passage without inducing significant deformation stress on the cells.

### 2.5 Assembly of the circulating flow system

The assembled chamber was connected to a reservoir containing culture medium and to a pump that provided continuous perfusion through the channel. The reservoir consisted of a Petri dish connected on both sides by tubing. This design enabled additional removal of air bubbles and stabilization of flow hydrodynamics. Culture medium or tumor cell suspension was delivered through the inlet port of the chamber and passed along the collagen gel containing fibroblasts. This architecture allowed modeling of intravasation and extravasation processes under controlled flow rates and reproducible mechanical conditions.

### 2.6 Mathematical modeling of the fluid flow in the chamber

To investigate velocity profiles and evaluate shear rates, a mathematical model of fluid motion in the channel of the flow chamber was developed. The computational domain was a rectangular parallelepiped with fixed length (4 cm) and height (2 mm) (see Fig. 6A, B). The chamber width W was varied in the simulations and assigned values of 3, 4, and 5 mm. A viscous (viscosity μ = 0.94 cP [12]), incompressible Newtonian fluid (density ρ = 1 g/cm^3^) entered the computational domain through a circular inlet on the parallelepiped wall, where a constant-pressure boundary condition was imposed. The fluid exited the chamber through another circular outlet on the same wall, where a constant mass flow rate Q was applied. Q was set to 0.5 mL/min, 1 mL/min, and 1.5 mL/min, values close to experimental conditions. Within the computational domain, the steady-state Navier–Stokes equation was solved numerically: ρ(u∇)u = μΔu − ∇p, where u is the fluid velocity and p is pressure. Heat-maps of shear rates and linear velocities were generated in the cross-sections shown in Fig. 6A, B. Hydrodynamic calculations were performed using the FlowVision software package (TESIS, Moscow, Russia) [13]. Graphs were produced in OriginPro (OriginLab Corporation, Northampton, Massachusetts, USA), and figures were prepared in Adobe Illustrator (Adobe, San Jose, California, USA).

## 3. Results

A biotechnological flow-based system designed for studying tumor cell intravasation and extravasation was developed and experimentally validated. In the first stage, the optimal collagen gel concentration was selected to ensure matrix stability, cell viability, and sufficient mechanical robustness for forming a channel between the plates. In the second stage, the feasibility of co-culturing different cell populations within the gel was tested, including human dermal fibroblasts embedded in collagen and the endothelial cell line Ea.hy926 on the gel surface. In the third stage, the flow chamber was assembled and tested. In the fourth stage of the study, a mathematical model was constructed to determine the range of shear rates generated in the final chamber design under varying channel widths and volumetric flow rates of fluid perfused through the system.

### 3.1 Selection of the collagen concentration

At the first stage of the experiment, a series of tests was conducted to determine the optimal concentration of the collagen gel. Three collagen concentrations (10, 20, and 30 mg/mL) were compared based on mechanical stability, morphology, viability of encapsulated cells, and stability during chamber assembly. The latter two parameters were assessed using light microscopy and staining (Fig. 2).

**Figure 2.**
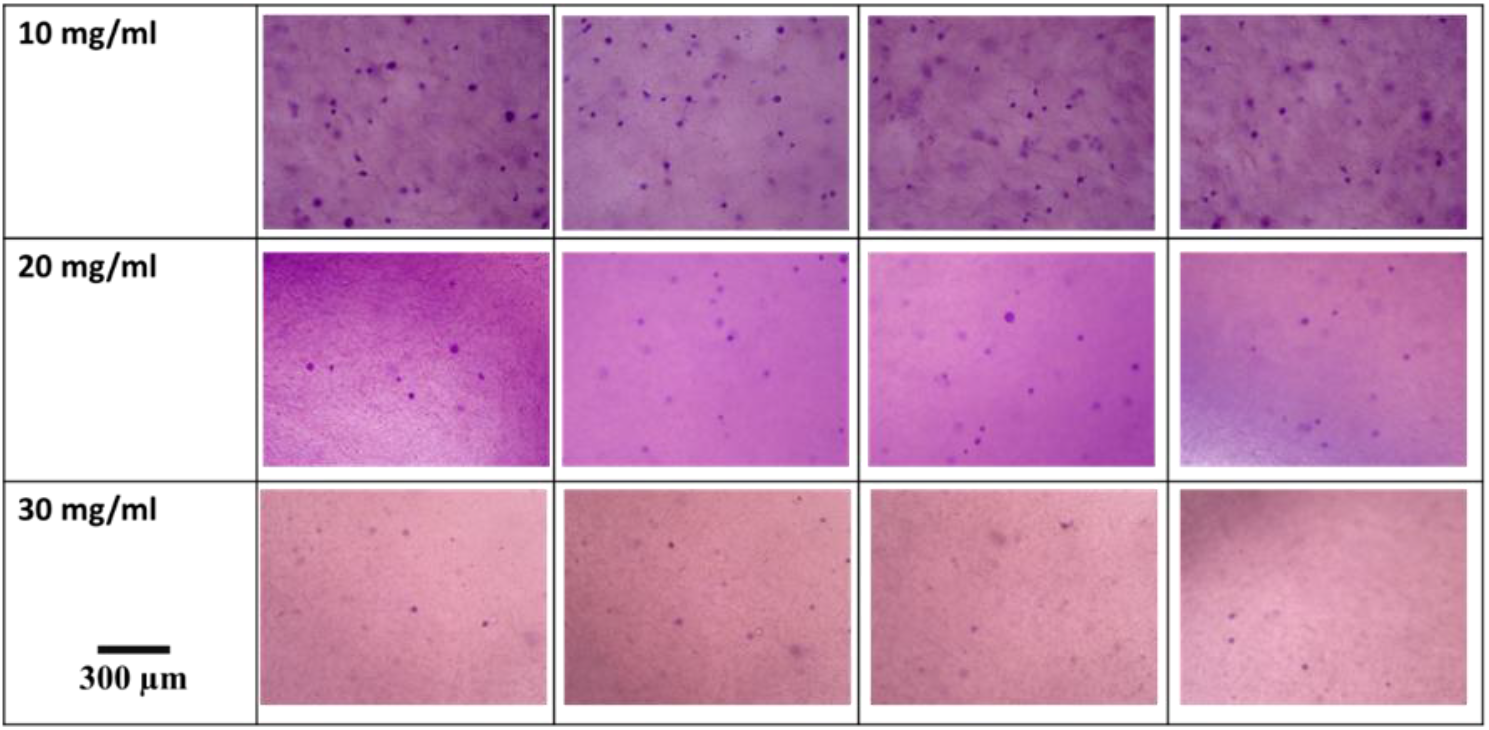
Light microscopy illustrating the effect of gel concentration on the morphology of human dermal fibroblasts.

The gel with a concentration of 10 mg/mL was characterized by low density and high deformability under any mechanical manipulation. When applied to a substrate, it spread immediately and did not retain its shape. This would make it unsuitable for creating a channel during assembly of the flow system. Visualization of fibroblasts within this gel showed good distribution and morphology. Gels with a concentration of 20 mg/mL demonstrated the optimal balance of mechanical stability and cell compatibility. The matrix retained its shape during handling and withstood mechanical load. Fibroblasts were evenly distributed throughout the gel and maintained normal morphology. The Hematoxylin/Eosin staining revealed well-distinguishable cells with an elongated fibroblast-like shape. At a concentration of 30 mg/mL, the gel exhibited high density and stiffness, likely reducing the diffusion of nutrients and slowing cell growth. Morphologically, cells appeared to be more spherical. The gels were stable during manipulation but were less suitable for modeling cell–matrix interactions and hindered visualization of cells using light microscopy. Thus, a concentration of 20 mg/mL was selected as optimal for subsequent experiments, as it provided a combination of biomechanical reliability and favorable conditions for fibroblast viability.

### 3.2 Co-culture of cell lines

To evaluate the feasibility of forming a multicomponent microenvironment, the endothelial cell line Ea.hy926 was added onto collagen gel (20 mg/mL) containing fibroblasts. Endothelial cells were applied to the surface of the preformed gel after one day of incubation. Over the course of 8 days, light microscopy revealed good adhesion of Ea.hy926 cells to the collagen matrix surface and the formation of a cellular layer (Fig. 3).

**Figure 3.**
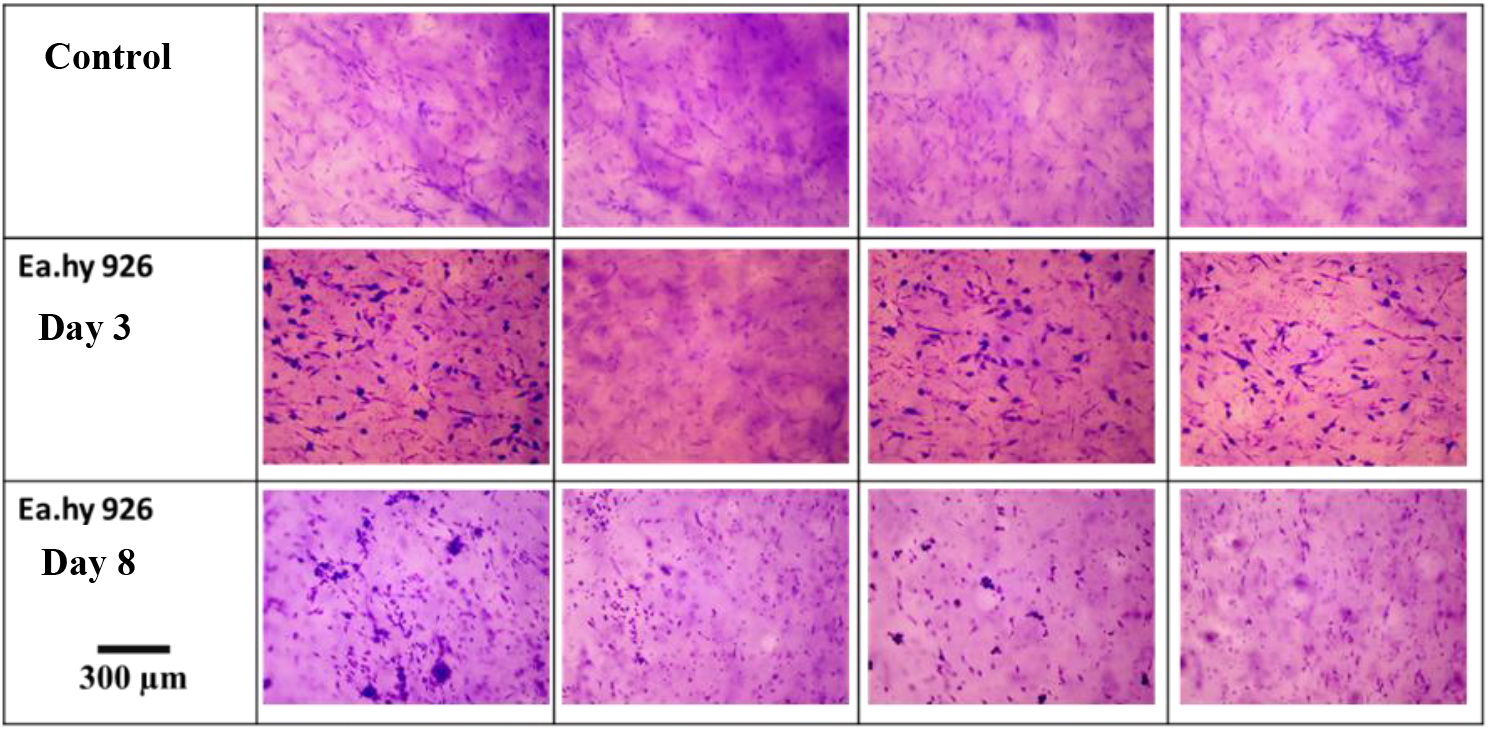
Collagen gel with human dermal fibroblasts (20 mg/mL); Ea.hy926 cells applied to the surface after one day of cultivation.

Fibroblasts embedded within the gel retained their morphology and viability, while the endothelial monolayer formed a surface covering. This confirms that the model can be used for studying interactions between tumor cells, stroma, and endothelium. Thus, the feasibility of creating a two-component cellular system—stromal cells within the matrix and an endothelial coating on its surface was demonstrated.

### 3.3 Development of the flow system

At the next stage, a flow chamber was assembled from three polyethylene terephthalate plates. The middle plate contained a central cavity for introducing collagen gel. The upper plate had inlet and outlet openings spaced 30 mm apart. The thickness of the channel between the plates was 2 mm. The plates were joined using sterile double-sided adhesive tape, ensuring structural integrity and leak-proof assembly. The completed chamber was connected to a reservoir containing culture medium and to a pump providing continuous perfusion through the channel (Fig. 4).

**Figure 4.**
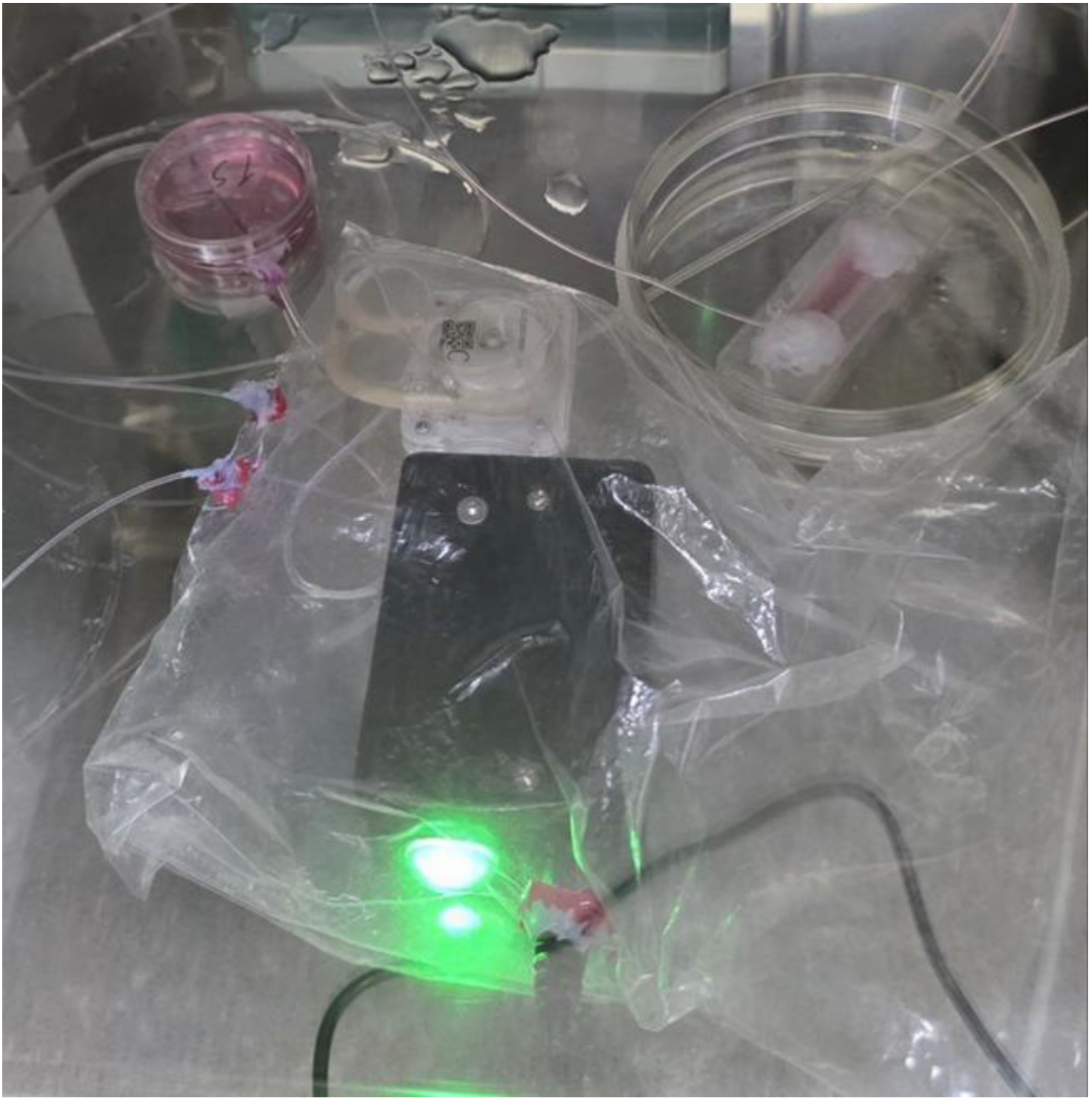
Assembled configuration of the system.

Testing of the system demonstrated that the 20 mg/mL collagen gel formed a stable channel that maintained its geometry over several days. Perfusion of full medium via the pump ensured a consistent flow. The additional open reservoir effectively removed air bubbles, stabilizing the flow and preventing pressure fluctuations. The system withstood prolonged medium circulation and was suitable for experiments involving suspensions of tumor cells. Fibroblasts within the gel remained viable even on day 8 of perfusion, confirming both the biocompatibility and the structural stability of the system (Fig. 5).

**Figure 5.**
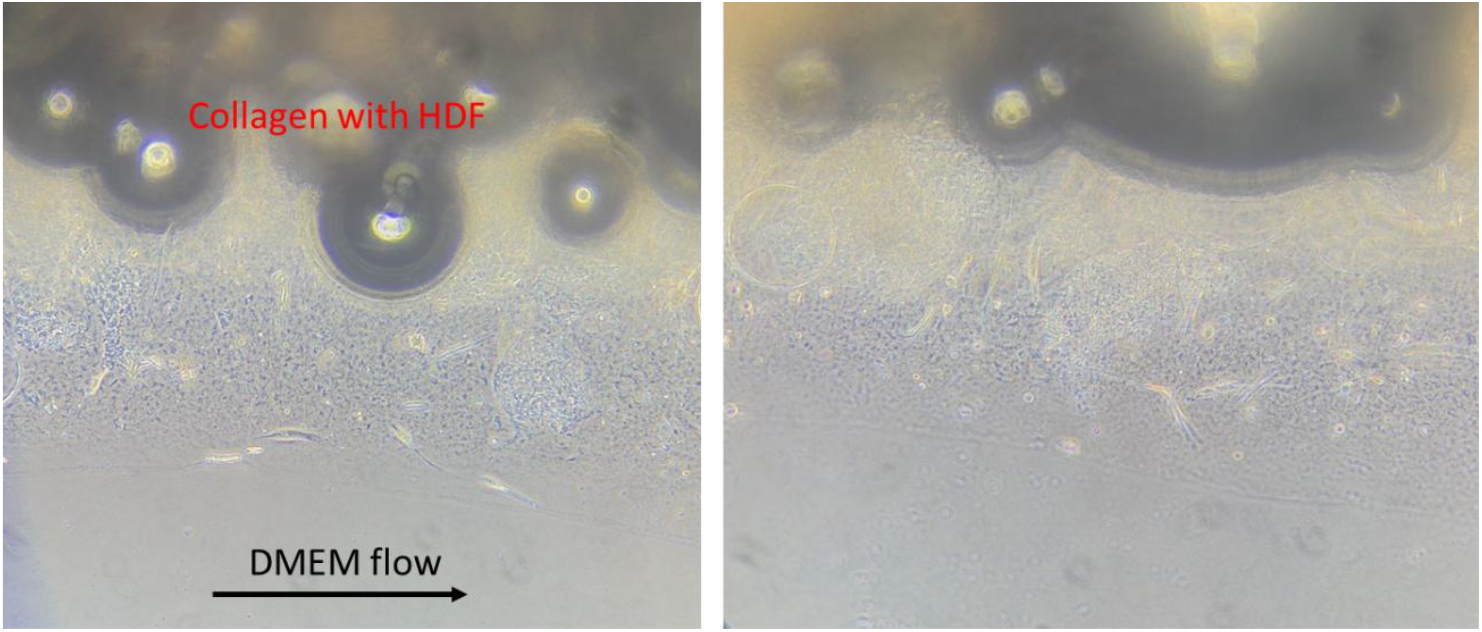
Light microscopy image of the chamber containing human dermal fibroblasts on day 8 of incubation under continuous perfusion with complete medium.

### 3.4 Mathematical modeling of flow velocity in the chamber

The characteristic distributions of linear velocity fields (Fig. 6C, D, E, G, H) and shear rate fields (Fig. 6 F, I, J) are similar for all chamber widths W and mass flow rates Q. The highest linear velocities (up to 400 mm/s) occur in close proximity to the inlet and outlet of the flow chamber (Fig. 6C, E). Fluid moves between these two points with characteristic velocity of approximately 3 mm/s (Fig. 6 D, G, H). In the regions between the chamber side faces and the nearest inlet/outlet, velocities are even lower than in the central part of the chamber. Shear rates range from 104 s^−1^ near the inlet/outlet to zero in the center of the chamber (Fig. 6 F, I, J). The highest shear rates are observed at the inlet and outlet. These elevated shear values remain within a distance of no more than 0.3 mm from the inlet/outlet, as illustrated for a chamber width of 4 mm at a flow rate of 1 mL/min (Fig. 6F). The regions with the greatest shear correspond to the edges of the inlet (diameter 0.3 mm). Shear rates along the remaining chamber walls, located far enough from the inlet and outlet, are of the order of 100 s^−1^ (Fig. 7). Moving from the walls toward the center of the chamber, shear rate decreases. In the following analysis, the term “near-wall shear rate” will refer to the maximum shear value measured across the longitudinal-section shown in Fig. 6J for each simulation. This cross-section passes through the center of the chamber and does not include regions near the inlet or outlet where extreme shear values (104 s^−1^) occur. An increase in volumetric flow rate in chambers of any width leads to a corresponding increase in near-wall shear rate (Fig. 7). Similarly, decreasing the chamber width results in an increase in shear rate (Fig. 7).

**Figure 6.**
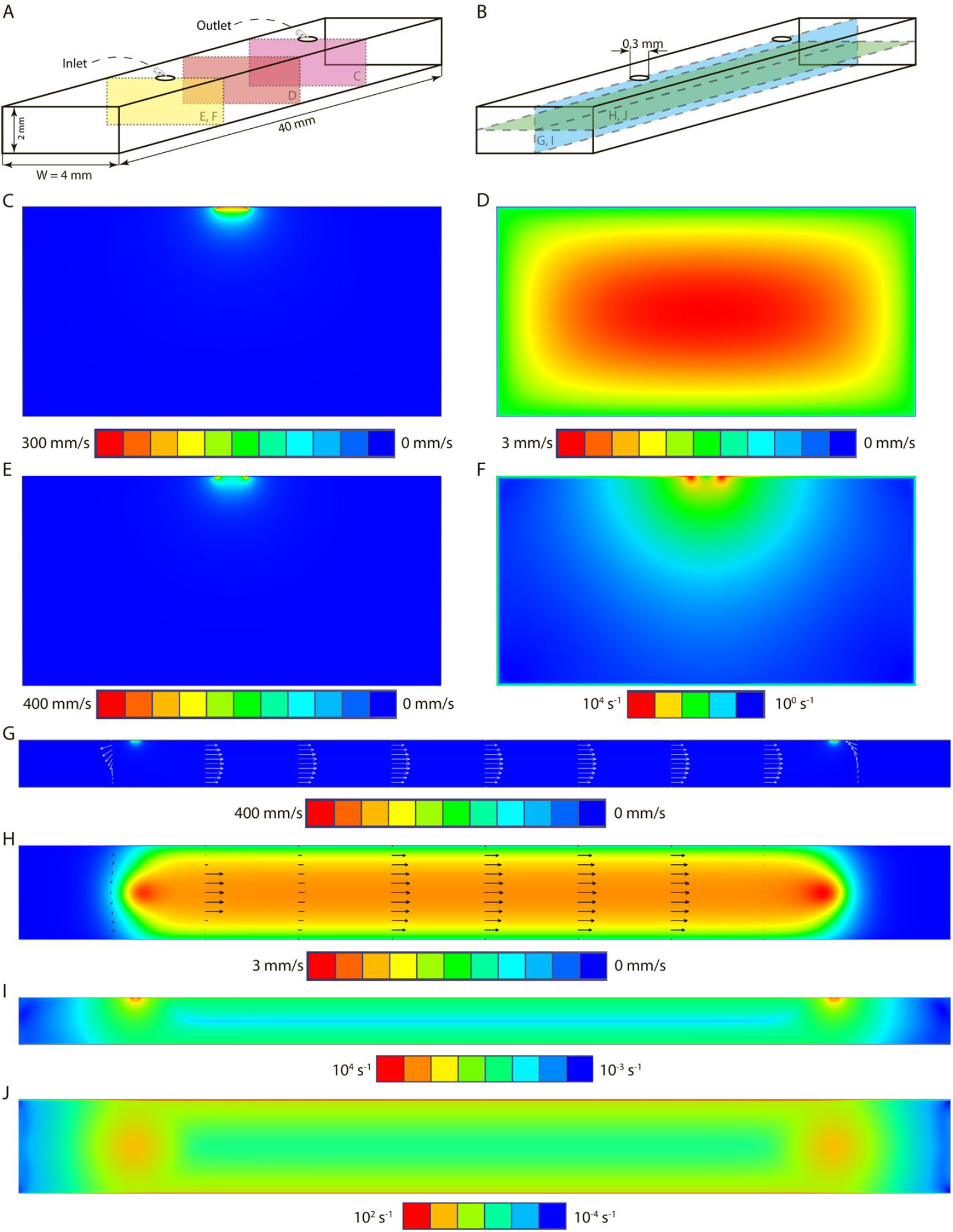
Scheme of the computational domain and results of the modeling for the flow chamber shown here for a chamber of width W = 4 mm and flow rate Q = 1 mL/min. A) Scheme of the flow chamber with depicted dimensions, including the width W, which was varied through the simulations. Colored regions denote position of cross-sections in which the velocity and shear-rate fields are shown. Cross-sections are labeled with letters corresponding to the panels in Fig. B) Diagram of the flow chamber with colored longitudinal-sections. Color scales are linear for linear velocity fields and logarithmic for shear-rate fields. C) Linear velocity field in the cross-section passing through the center of the outlet. D) Linear velocity field in the cross-section passing through the center of the chamber. E) Linear velocity field in the cross-section passing through the center of the inlet. F) Shear rate field in the cross-section passing through the center of the inlet. The color scale is logarithmic. G) Linear velocity field in the longitudinal section passing through both openings; arrows indicate flow direction. H) Linear velocity field in the longitudinal section parallel to the plane containing both openings; arrows indicate flow direction. I) Shear rate field in the longitudinal section passing through both openings. J) Shear-rate field in the longitudinal section parallel to the plane containing both openings.

**Figure 7.**
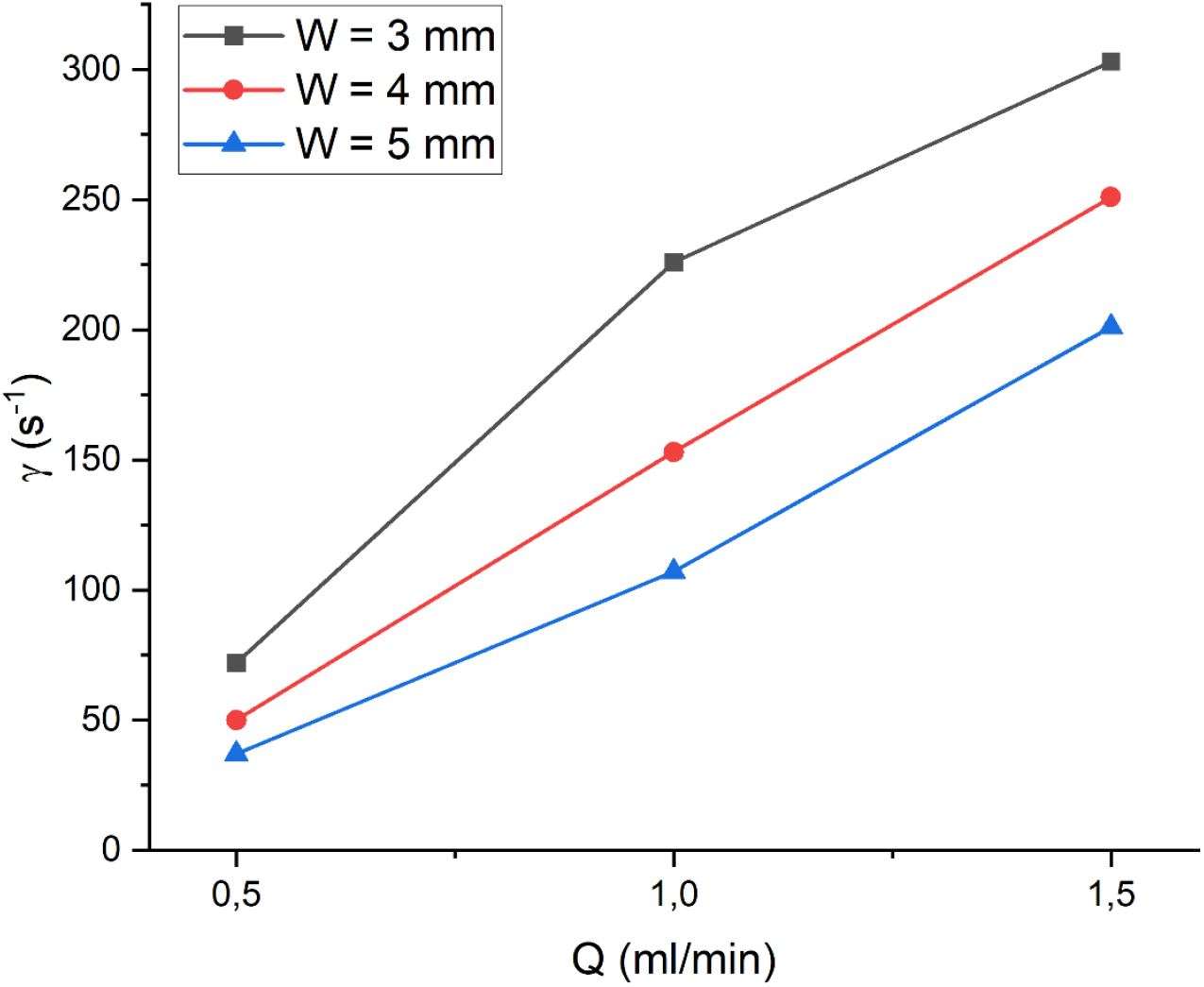
Dependence of the wall shear rate in the flow chamber on the mass flow rate of fluid passing through the chamber. Wall shear rate was defined as the maximum shear-rate value across the cross-section shown in Fig. 6K for each simulation.

## 4. Discussion

The flow-based biotechnological system developed in this study reproduces key elements of the microenvironment that govern tumor cell intravasation and extravasation. The obtained results demonstrate that the proposed model is capable of addressing limitations inherent to existing 2D, 3D, and microfluidic platforms, which insufficiently capture the biophysical characteristics of vascular flow and cell–extracellular matrix interactions. It is well established that mechanical and spatial cues determine whether tumor cells can penetrate into or exit from a vessel; however, most experimental systems do not provide a combination of perfusion, a 3D matrix, and physiologically relevant channel dimensions [1,2].

The optimal collagen concentration of 20 mg/mL identified in our study provided matrix stability, uniform fibroblast distribution, and long-term preservation of cell viability. Experimentally, it was observed that gels that were too soft deformed under mechanical load, whereas overly dense matrices affected cellular behavior and hindered visualization by light microscopy. Thus, the use of 20 mg/mL is well justified.

The addition of endothelial Ea.hy926 cells to the gel surface enabled the formation of a two-component microenvironment combining a 3D stromal compartment with an endothelial barrier. The observed development of a cohesive endothelial layer over several days confirms the suitability of the system for modeling transendothelial migration of tumor cells—a process that is still studied predominantly in Transwell systems. Meanwhile, modern reviews emphasize the necessity of transitioning to dynamic models, since the behavior of both endothelium and tumor cells changes dramatically under the influence of flow, shear stress, and pressure [5,6].

A key advantage of the developed system is its ability to maintain a stable flow without bubble formation and without deformation of the channel. Most microfluidic devices are limited by small channel dimensions (50–150 μm), which induce mechanical deformation of circulating tumor cells, activate RhoA/ROCK-dependent stress pathways, and distort the physiological mechanisms of intravasation and extravasation [8]. In contrast, the 2-mm-high channel created in our system does not impose excessive compressive stress on cells and more closely approximates physiological conditions.

Stabilization of pressure and flow using an additional open reservoir represents an important engineering solution. This design feature prevents bubble formation—one of the most frequent problems in microfluidic systems, where bubbles disrupt laminar flow, damage the endothelium, and lead to non-representative data [9]. In our system, this limitation is effectively eliminated, ensuring reproducibility during long-term experiments.

An additional advantage is the possibility of using large millimeter-scale gels, which surpasses what can be achieved in microfluidic platforms. The matrix thickness in such systems typically ranges from 100 to 300 μm and cannot reproduce the physiological diffusion and mechanical gradients characteristic of the tumor microenvironment [10]. Our data demonstrate that fibroblasts maintain their viability and morphology even during prolonged perfusion, confirming the appropriateness of the selected gel composition and the efficiency of the flow system.

Results of computational modeling showed that the shear rates generated in the developed flow chamber are close to 100 s^−1^ across the full range of channel widths formed by the collagen gel layers. This corresponds to shear conditions in human venous circulation. By adjusting channel width and pump rate, wall shear rates could be tuned from tens to hundreds of reciprocal seconds. However, at short distances (less than the diameter of the inlet/outlet) from the inlet and outlet, shear rates reach 10^4^ s^−1^, which must be taken into account when analyzing the state of components circulating within the chamber.

Thus, comparison of the obtained results with existing literature allows us to conclude that the developed model integrates the advantages of 3D hydrogels, dynamic bioreactors, and microfluidic systems while avoiding their major technical limitations. As highlighted in recent reviews, physiologically relevant geometry and the ability to work with real biological fluids are critical parameters for studying the metastatic cascade [6,7]. Our system meets these criteria, making it a promising tool for investigating mechanisms of intravasation and extravasation, tumor–stroma interactions, and for testing anti-metastatic therapies.

## 5. Conclusion

In this study, a flow-based biotechnological system designed for investigating tumor cell intravasation and extravasation under controlled perfusion was developed and experimentally validated. An optimal collagen gel concentration (20 mg/mL) was identified, providing mechanical stability of the matrix and long-term fibroblast viability. The feasibility of applying endothelial cells onto the gel surface was successfully demonstrated, enabling the formation of a cellular barrier that mimics the vascular wall. The chamber design provided stable flow without bubble formation due to the incorporated reservoir, representing a significant advantage over microfluidic models. The large size of the system also enables the use of native biological fluids and tumor cell suspensions without inducing deformation or stress. Using the mathematical model constructed for this chamber, estimates of the hemodynamic conditions inside the system were obtained. The results confirm that the developed system is a reliable, reproducible, and functional platform for studying the early stages of the metastatic cascade.

